# Split *Staphylococcus aureus* prime editor for AAV delivery

**DOI:** 10.1101/2021.01.11.426237

**Authors:** Eric J. Aird, Alina C. Zdechlik, Brian L. Ruis, Colette B. Rogers, Andrew L. Lemmex, Andrew T. Nelson, Eric A. Hendrickson, Daniel Schmidt, Wendy R. Gordon

## Abstract

Prime editing brings immense promise to correct a large number of human pathogenic mutations and enact diverse edit types without introducing widespread undesired editing events. Delivery of prime editors in vivo would enable such edits to be introduced in a clinical setting. The coding sequence for prime editor, however, is too large to fit within the size-constrained adeno-associated virus (AAV) genome. Herein, we describe a split *Staphylococcus aureus* prime editor capable of being delivered by dual AAVs. We characterize the editing ability of plasmid-based versions of an *S. aureus* prime editor in vitro at a variety of loci with diverse edit types. We investigate various split prime editor architectures and alternative dimerization domains. Finally, we demonstrate the capacity of prime editor to be co-delivered by dual AAVs in vitro. While editing rates are lower than desired, this approach presents an important step to translate prime editing for in vivo delivery.

## Introduction

Genome editing has brought incredible promise to correct or ameliorate previously untreatable genetically linked diseases such as Tay-Sachs disease or Phenylketonuria (PKU). The use of programmable nucleases such as CRISPR-Cas9 that introduce a DNA double-stranded break (DSB) enables precise gene editing at a desired locus^1,2^. Traditionally, homology-directed repair (HDR) has been the only means to introduce precise nucleotide changes at a DSB via the supplementation of a donor DNA molecule encoding the changes. HDR, however, usually occurs less frequently than the circumstantially undesired non-homologous end joining (NHEJ) pathway^3^. To circumvent the NHEJ pathway, base editors were designed to change single bases in DNA without the need to create a DSB^4^. Base editors employ nickase Cas9s that have one of the nuclease domains mutated to prevent cleavage on one DNA strand. Base editors, as the name suggests, are limited to introducing transition point mutations (purine:purine or pyrimidine:pyrimidine), although recently C to G base editors have been characterized^5,6^. While NHEJ levels are low with base editors and high rates of precise editing can be achieved, the restraint on editing type and inability to edit specific bases within a window makes it not ideal for many desired precise genome editing applications.

The latest advent in CRISPR genome editing technologies is prime editor, which is capable of introducing a wide change of precise modifications without the need for DSB formation^7^. Like base editor, prime editor utilizes a Cas9 (H840A) nickase. A reverse transcriptase (RT) is tethered to the nickase while a prime editing guide RNA (pegRNA) contains a 3’ extension serving as both the genome hybridizing site (primer binding site; PBS) and the RT template encoding the desired edits. Prime editor cleaves the single strand of DNA, the PBS of the pegRNA hybridizes with the newly exposed ssDNA, and the RT synthesizes from the RT template. Flap resolution and DNA mismatch repair then allow for incorporation of the desired modification. Therefore, prime editing is theoretically able to make edits downstream of the nick site, termed the +1 site. Prime editor was shown to allow for edits ranging from codon changes to small insertions and deletions through encoding these modifications in the 3’ pegRNA extension^7^. Hypothetically, prime editor is capable of correcting close to 90% of human pathogenic mutations^7^. As with other types of genome editing technologies, however, in vivo targeting and delivery remains a large hurdle to overcome to achieve such a goal^8^.

Adeno-associated virus (AAV) is the most common in vivo delivery vehicle for gene editing reagents. AAV contains a single-stranded DNA (ssDNA) genome that can be replaced with transgenes of interest. The virus can target a wide range of tissue and cell types with a relatively low immunogenicity profile. AAV-mediated delivery of CRISPR-Cas has been successfully deployed to produce indels^9^, create chimeric T-cell receptors^10^, and generate large deletions^11^. Extensive engineering efforts have also modified the virus to enhance cell-specific delivery or decrease immunogenicity^12^. A major constraint of AAV is the maximum size of its ssDNA genome, roughly 4.7 kb not including the two flanking inverted terminal repeats (ITRs)^13^. The conventional Cas9 originating from *S. pyogenes* (SpCas9) is 4.2 kb in size, allowing little space for necessary regulatory elements such as promoters and terminators and precludes encoding of a gRNA expression cassette in the same genome. To overcome this limitation, smaller orthologs of Cas9 such as from *S. aureus* (SaCas9) have been utilized^14^. The gene length of SaCas9 is approximately 1 kb shorter than SpCas9 allowing for more freedom in packaging in AAV. Another strategy to deliver full-length Cas proteins has been to split the protein into two AAV vectors to be co-delivered. This approach can take place on the DNA, RNA, or protein level (reviewed in ^15^). On the protein level, engineered split trans-splicing inteins can be co-opted essentially as dimerization domains to bring two halves of a split protein together^16^. For base editor to be packaged and delivered by AAV, a split base editor, split intein system was used^17,18^. The two intein halves associate at low nM affinity and swiftly excise themselves out, leaving only a small scar. For both SpCas9 and SaCas9, various groups have identified numerous permissible split locations that allow the proteins to retain nearly full activity once recombined^18,19^.

Prime editor was previously shown to be delivered ex vivo using a three-part lentivirus system to mouse primary cortical neurons^7^. This approach, however, is not broadly applicable in the clinic due to constraints on choice of cell type to edit. Ribonucleoprotein (RNP)-containing lentiviral particles have been used to transiently deliver Cas9^20^, but the sheer size of prime editor protein (∼250 kDa) makes it difficult to produce conventionally in *E. coli* and package. Ideally, AAV would be used to deliver prime editor. At ∼6.4 kb, prime editor is too large to be encoded into a single AAV genome. Even when constituted as a split version, prime editor is unable to fit into two AAV genomes with the necessary regulatory elements and the pegRNA expression cassette.

To address this size constraint issue, we developed a new version of prime editor using SaCas9(N580A) as the guided nickase. We demonstrate that *S. aureus* prime editor (SaPE) can also introduce targeted changes to a wide range of genomic loci from point mutations to larger deletions. We assessed three different split locations in concert with split intein and split NanoLuc approaches to reconstruct full-length, active protein. All versions of split SaPE were capable of inducing editing events, although they tended to be less efficient than the full-length protein. The split version can be packaged into two standard size AAV genomes and be delivered in viral form in vitro. This work opens a potential avenue for in vivo and clinical exploration of prime editing.

## Results

### *Development of an* S. aureus *prime editor*

As SaCas9 and SpCas9 share similar domain architecture, we reasoned that a nickase SaCas9 could serve as a sufficient nicking nuclease to be combined with the prime editing methodology. We generated the analogous HNH domain mutation in SaCas9 (N580A) as in SpCas9 (H840A) and tethered a reverse transcriptase to create an *S. aureus* prime editor, SaPE (**Figure 1a**). The previously engineered version of Moloney murine leukemia virus reverse transcriptase (eMMLV-RT) was utilized^7^. We then sought to create a split version of SaPE capable of fitting into two AAV genomes. Numerous methods have been applied to efficiently recombine split portions of Cas9^21,22^. We utilized a split intein-mediated approach wherein the split Npu trans-splicing intein from *Nostoc punctiforme* was appended onto the N- and C-termini of split SaPE^23^ (**Figure 1a**). We trialed three different split locations in SaCas9, all of which have been previously reported^18,19^. Two of the split locations, E739/S740 (version 1) and K534/C535 (version 3), showed reasonable intein splicing in cells (**Figure 1b**).

**Figure 1.**
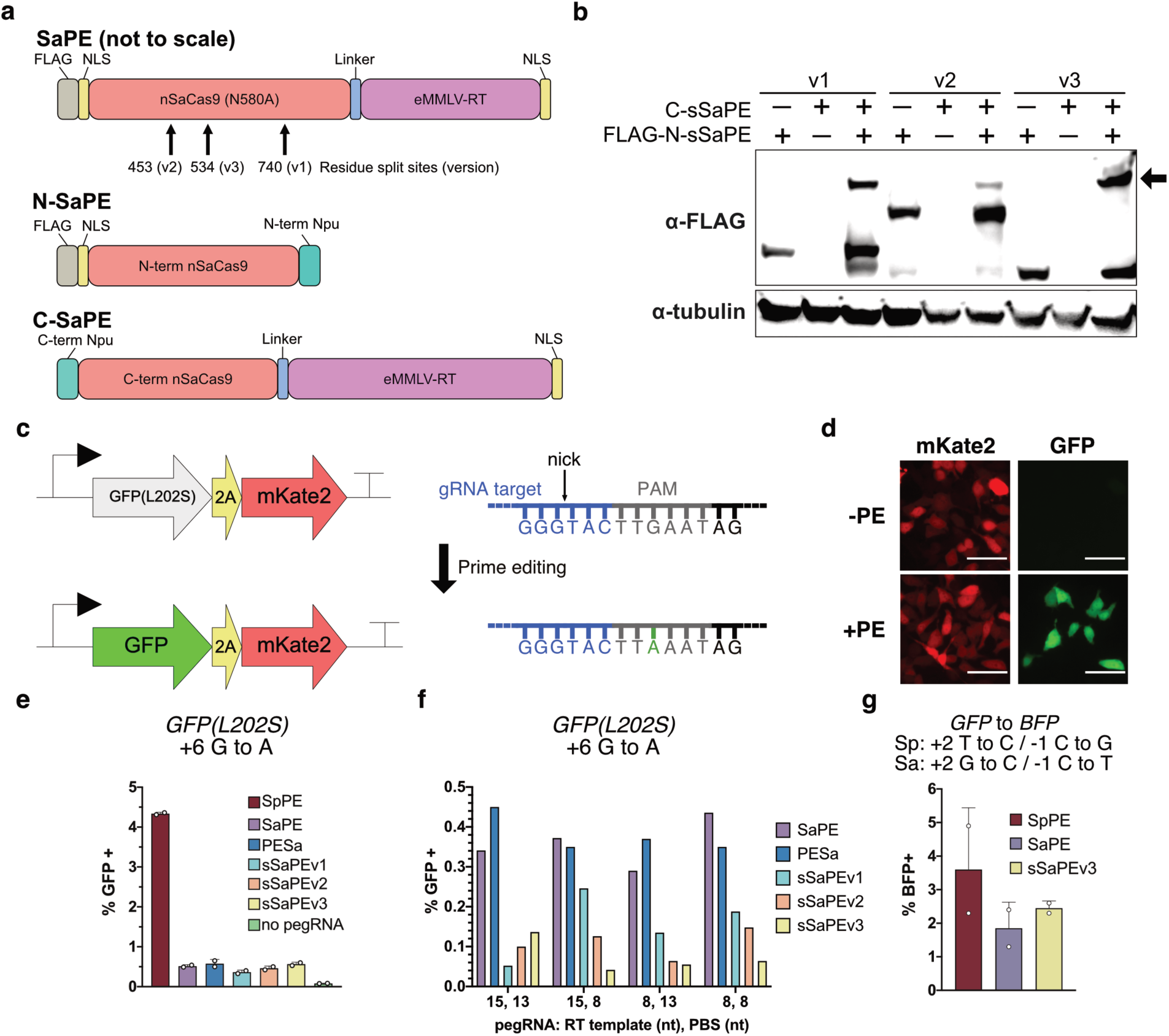
Enabling precise genome modifications using a split *S. aureus* prime editor. (**a**) Schematic of both full-length *S. aureus* prime editor (SaPE) and split SaPE (sSaPE). (**b**) Western blot of plasmid-based expression of three different versions of sSaPE. Successful trans intein splicing is denoted by the arrow next to the higher molecular weight band. (**c**) Model system for analyzing editing by SaPE using a GFP(L202S) stable reporter line to restore GFP fluorescence. A single point mutation (G to A) that lies within SaPE’s PAM is required. The antisense DNA sequence is shown. (**d**) Representative fluorescence microscopy images of unedited (-PE) and edited (+PE) HEK-293T stably expressing the GFP(L202S)-2A-mKate2 reporter. Scale bar = 20 µm. (**e**) Quantitation of prime editing in HEK-293T cells using flow cytometry. *S. pyogenes* prime editor (SpPE) is compared to full-length SaPE, N-terminally fused RT-nSaCas9 (PESa), and three versions (v1-v3) of sSaPE. No pegRNA corresponds to transfection of SaPE only. Data are representative of multiple independent experiments. (**f**) Targeting of *GFP(L202S)* reversion with pegRNAs of differing 3’ extension lengths (PBS = primer binding site). (**g**) GFP to BFP editing in HEK-293 cells stably expressing wildtype GFP. Individual data points represent biological replicates.

To more quickly test editing conditions, we created a stably integrated fluorescent reporter cell containing a point mutation in *GFP(L202S)* that ablates GFP fluorescence linked to mKate2 via a 2A self-cleaving peptide^24^ (**Figure 1c**). The restoration of GFP fluorescence through a +6 G to A transition point mutation can readily be detected using microscopy or flow cytometry (**Figure 1d**). We first tested full-length and split versions of SaPE alongside a variant with the RT tethered to the amino terminus of nSaCas9 (PESa). In comparison with SpPE, SaPE had an 8-fold reduction in editing efficiency (**Figure 1e**). While SaPE editing efficiencies were low at around 0.5%, all split versions had comparable levels to full-length SaPE. These low editing frequencies with SaPE, as elaborated on in the Discussion, are a trend that generally holds true at most loci. Decreases in editing efficiency might be attributed to, among other factors, shorter residency time of SaPE on the DNA as opposed to SpPE^25^. Nonetheless, detectable levels of prime editing driven by SaPE are observed.

Prime editing necessitates optimization of the pegRNA 3’ extension design as both the RT template and primer binding site (PBS) lengths appear to be edit type and locus dependent^7,26,27^. At the *GFP(L202S)* locus, we interestingly observed little difference in editing frequencies when sampling altered 3’ extension designs (**Figure 1f**). Variation did exist among the three split versions, and in these sets of experiments, split SaPE had noticeable decreased editing frequencies compared to full-length. Combined with the intein splicing and editing profiles, we carried forward predominantly with version 3 of sSaPE (K534/C535).

In another fluorescent assay targeting a different locus in *GFP*, we once again compared SpPE to SaPE. A two amino acid change in GFP (T65S-Y66H) converts the fluorescence to BFP^28^. When prime editing reagents were transfected in HEK-293 cells stably expressing GFP, we observed a similar frequency of editing between full-length and split SaPE (1.85% versus 2.45%) (**Figure 1g**). This was again lower than SpPE (3.6%) but to a lesser degree than *GFP(L202S)* where SpPE editing was almost an order of magnitude higher. Conversion to BFP was also seen in HCT-116 cells to a similar degree as in HEK-293 cells (**Supplementary Figure 1**). Interestingly, PAM sequence constraints required this edit to be upstream from the nick site (−1 site) in addition to a +2 point mutation. The ability of prime editor to make modifications upstream of the predicted nick site, 3 bp away from PAM, lends itself to past research suggesting Cas9 also can cleave 4 bp upstream^29^. While this effect might be locus dependent, we observed appreciable editing and potentially a broadened capability of prime editor.

### Optimization of split SaPE

To try increasing editing frequencies at the *GFP(L202S)* locus with SaPE, we next sought to optimize experimental parameters and platform design. We performed a titration of plasmid DNA concentration using lipofection while the molar ratio of PE:pegRNA was held constant. Due to the PE vectors being approximately three times the size of the pegRNA plasmids in bp, this resulted in a 3:1 ng of DNA concentration ratio. The maximal editing of ∼0.6% was seen at surprisingly the lowest concentration of DNA. (**Supplementary Figure 2**). In the case of sSaPEv3, higher DNA concentrations resulted in roughly a 33% decrease in editing efficiency. Cytotoxicity associated with large plasmids combined with excess amounts of DNA could be a cause of this trend^30^.

Next, two previously described linkers were installed in place of the built-in one between nSaCas9 and RT to try enhancing editing rates: XTEN^31^, found in base editors, and (H4)_2_, a rigid alpha helical linker of the sequence [A(EAAAK)_4_A]_2_ ^32^. While the XTEN linker performed as well as the original SaPE linker, the replacement with the (H4)_2_ linker ablated prime editing (**Supplementary Figure 2**). This difference highlights the crucial, but often ignored, impact of spatial orientation of components in fusion proteins. Further exploration of linkers and different permissible fusion locations of RT on nCas9 could be carried out to try optimizing this key parameter.

Another approach we utilized to try boosting editing rates was to employ the 3^rd^ generation prime editor system, termed PE3^7^. PE3 uses a 2^nd^ gRNA that nicks the non-prime edited strand to encourage DNA repair machinery to preferentially repair the nicked strand once the desired edit has been incorporated. The hypothesis is to push the equilibrium of flap resolution and DNA repair towards the desired outcome, a method successfully used in base editors^4^. Due to the limited number of SaCas9 PAM motifs in the vicinity of the targeted location in *GFP(L202S)*, we could only assess two PE3 gRNAs in combination with SaPE. Use of either gRNA, which nick at −70 or −51 bp from the +1 site, resulted in lower editing efficiencies for sSaPEv3 (**Supplementary Figure 2**). For sSaPEv1, the −51 gRNA caused a dramatic tripling in editing efficiency (0.20% to 0.67%) while the −70 gRNA had little effect. PE3 has been shown to be moderately successful at improving prime editing at other loci with SpPE^7,33^. We chose, however, not to generally pursue the PE3 approach at other loci due to the limited amount of neighboring PAM sequences available. Taken together, our efforts to optimize SaPE did little to improve editing rates from the outset in the context of the *GFP(L202S)* locus.

### Broad assessment of SaPE

We next wanted to assess SaPE and sSaPEv3 across a wide range of loci and editing types while also changing the lengths of the 3’ extension components of pegRNA. Using next-generation sequencing, we found SaPE capable of making wide-ranging precise changes to target loci (**Figure 2**). In sampling different RT template and PBS lengths at the *EMX1* locus in a +6 G to T transversion point mutation, we found variable editing rates between 0.3% and 2.1% for full-length SaPE (**Figure 2a**). For split SaPE, less variability existed, but editing rates were limited to between 0.3% and 0.5%. Split SaPE tended to have a decreased editing frequency across all loci tested, a trend that is not uncommon with split Cas9^21,34^ but is in contrast to what was observed with split base editor^18^. At both the *FANCF* and *DNMT3B* loci, editing rates peaked around 0.4% for a range of point mutations, insertions, and deletions (**Figure 2b and c**). Point mutations, however, tended to yield higher editing efficiencies than insertion or deletions. In varying the RT template length at *RUNX1*, the longer length (15 nt) yielded improved editing over the shorter RT template (10 nt), regardless of the location of the edit from the nick site (**Figure 2d**). In the case of the further +7 G to A mutation, a 10 nt RT template resulted in nearly undetectable levels of editing.

**Figure 2.**
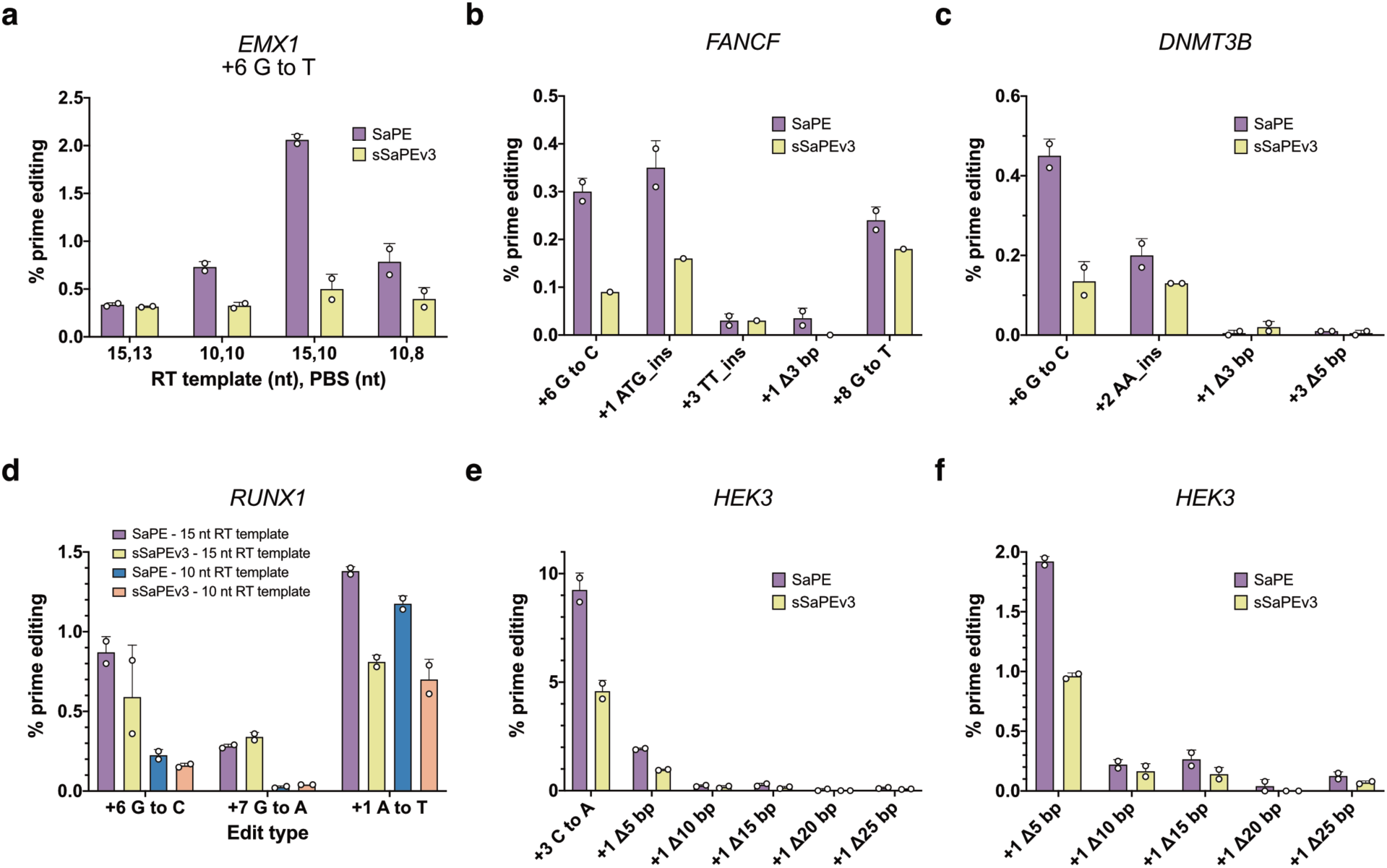
SaPE is capable of diverse edit types across various genomic loci. (**a-f**) Next-generation sequencing readouts of targeting of SaPE and sSaPEv3 to indicated loci (**a**) *EMX1*, (**b**) *FANCF*, (**c**) *DNMT3B*, (**d**) *RUNX1*, and (**e**,**f**) *HEK3* with differing edit types and pegRNA architectures. (**f**) contains the same 5mer deletion series data as (**e**), but with a scaled y-axis.

The highest prime editing frequencies were seen at the *HEK3* locus, where a +3 C to A transversion was formed in 9.2% of full-length SaPE transfected loci and nearly 5% in the sSaPEv3 condition (**Figure 2e**). At the same target site, a series of 5mer deletions were encoded in the pegRNA, ranging from 5 to 25 bp in length. Editing frequencies were 1.9% and 1% for SaPE and sSaPEv3, respectively, for a 5 bp deletion (**Figure 2f**). These rates decreased as the deletion length increased. 25 bp deletions were still detected, albeit at a lower frequency than other deletion lengths.

We repeated some of the above experiments in U2-OS cells to explore cell type dependency. Overall, no trend emerged in regard to predicting optimal editing frequency at a given target locus, edit type, and pegRNA design. Editing efficiencies were in the single digits across all loci assayed, although the *HEK3* locus offers promise moving forward.

### Altering the dimerization domain

Following extensive characterization of the editing frequencies, we aimed to better understand the split protein recombination requirements. To visualize reassociation of split SaPE, we replaced the Npu split intein with split NanoLuc (**Figure 3a**). Split NanoLuc (sNanoLuc, sNL), with a K_D_ of 700 pM, is comprised of an 18 kDa N-terminal fragment, LgBiT, and a 13 amino acid C-terminal portion, HiBiT^35^. When co-transfecting the two halves of sSaPE-sNanoLuc, we saw extensive nuclear reconstitution of NanoLuc in a vast majority of cells using bioluminescence microscopy (**Figure 3b**). Similar editing rates to sSaPEv3 were also seen, indicating that sNanoLuc is sufficient to act as a dimerization domain in the context of SaPE (**Figure 3c**). The lack of requiring covalent association of the two halves of SaPE was further evaluated by utilizing a catalytically inactive version of the Npu intein (C1A). In these constructs, the two intein components can associate but not self-splice. No covalent full-length SaPE is formed, yet editing rates are on par with the catalytically active intein version (**Figure 3c and d**). This data further validated that covalent association of the two halves of split SaPE is not required to reconstitute active SaPE. The use of sNanoLuc also offers a convenient system in which visual confirmation of recombined SaPE is possible, an approach which can be utilized with AAV delivery.

**Figure 3.**
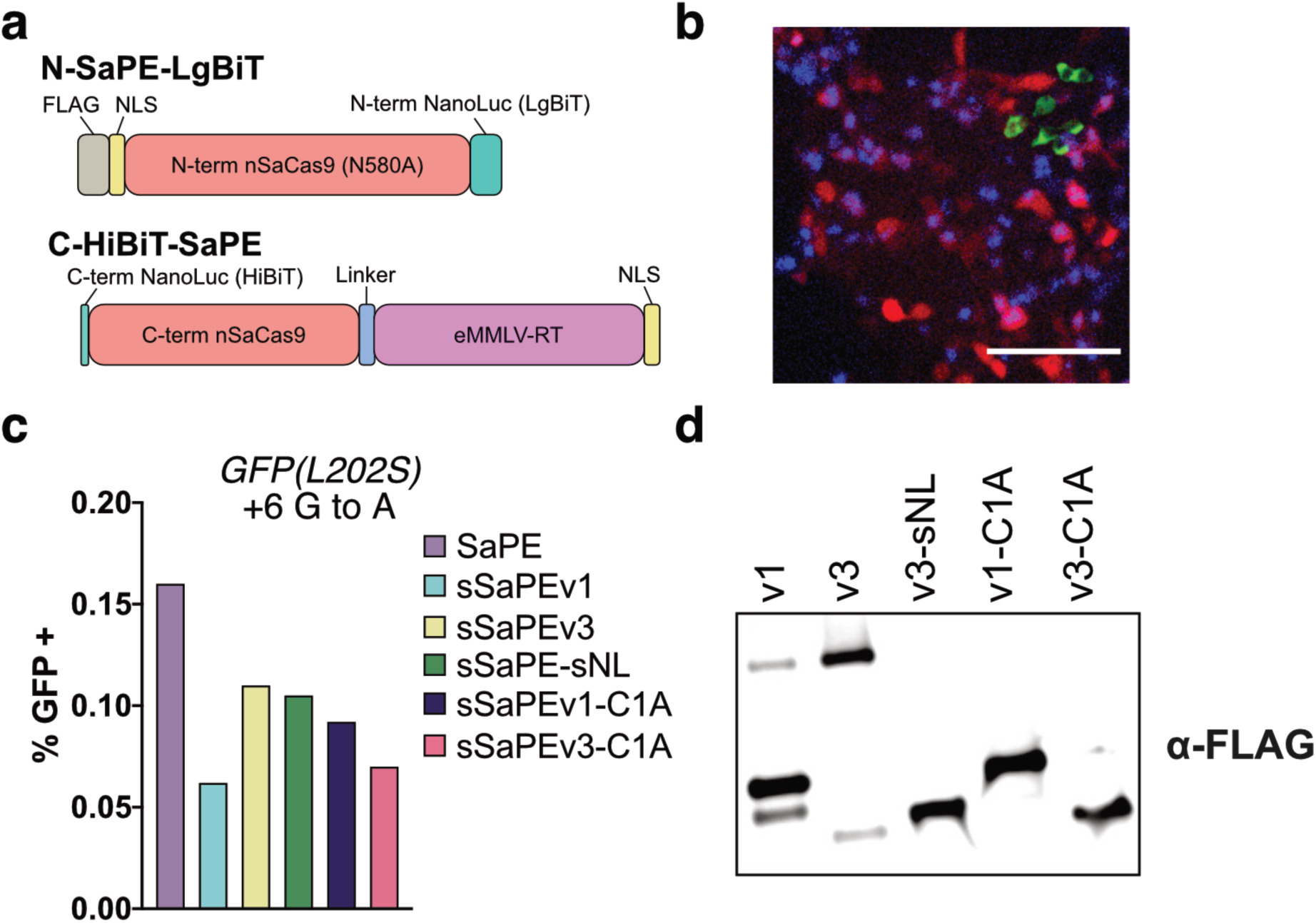
Split NanoLuc substitutes as an effective dimerization domain and visualization tool. (**a**) Diagram of sSaPE-split NanoLuc (sNL) (not to scale). (**b**) Composite image of stable GFP(L202S)-2A-mKate2 HEK-293T cells transfected with sSaPE-sNL. Colors are as follows: red = mKate2; blue = NanoLuc; green = GFP. Scale bar = 120 µm. (**c**) Prime editing rates at *GFP(L202S)* locus in HEK-293T cells. (**d**) Western blot of co-transfection of indicated versions of split SaPE. Only the N-terminal fragment is FLAG labeled. The top band corresponds to covalently recombined SaPE.

### Packaging and delivery of SaPE in AAV

Next, we cloned the N- and C-terminal portions of sSaPE versions 1 and 3 into AAV genomes containing flanking ITRs. For N-terminal sSaPE, two orientations of the U6 promoter-pegRNA cassette were tested, either in tandem or in reverse alignment to the protein expression cassette (**Figure 4a**). The C-sSaPEv3 AAV genome is 4.8 kb in length while the N-terminal genomes are approximately 3.5 kb. Before packaging into virus, AAV plasmids were co-transfected targeting *GFP(L202S)* to ensure editing still occurred in the different context. Indeed, editing rates were in line with previous plasmid designs, even with a co-transfection as opposed to a triple transfection (**Figure 4b**). Next, AAV-DJ was packaged and titered using qPCR, yielding ∼2×10^10^ viral genome copies (g.c.)/μl. A control transduction with tdTomato as the encoded genome exhibited robust delivery and fluorescent protein expression in HEK-293T cells (**Supplementary Figure 4**), ensuring proper production and purification of the virus. We co-transduced sSaPE AAVs in *GFP(L202S)* HEK-293T cells and first assessed protein recombination. At both three and five days post-transduction, full-length recombined protein was evident via western blot (**Figure 4c**). Assessing GFP fluorescence restoration in *GFP(L202S)*, we see a concentration dependent increase in editing (**Figure 4d**). While the GFP restoration frequency is low, co-delivery of sSaPEv3-sNanoLuc AAVs in these cells also showed a low number of cells expressing recombined protein, potentially due to a low co-transduction efficiency (**Supplementary Figure 4**). However, co-transduction in U2-OS cells resulted in an order of magnitude increase in delivery and recombination efficiency as visualized by bioluminescence microscopy (**Figure 4e**). Taken together, we have begun the proof of concept work necessary to allow prime editor to be delivered via dual AAVs for potential future clinical applications.

**Figure 4.**
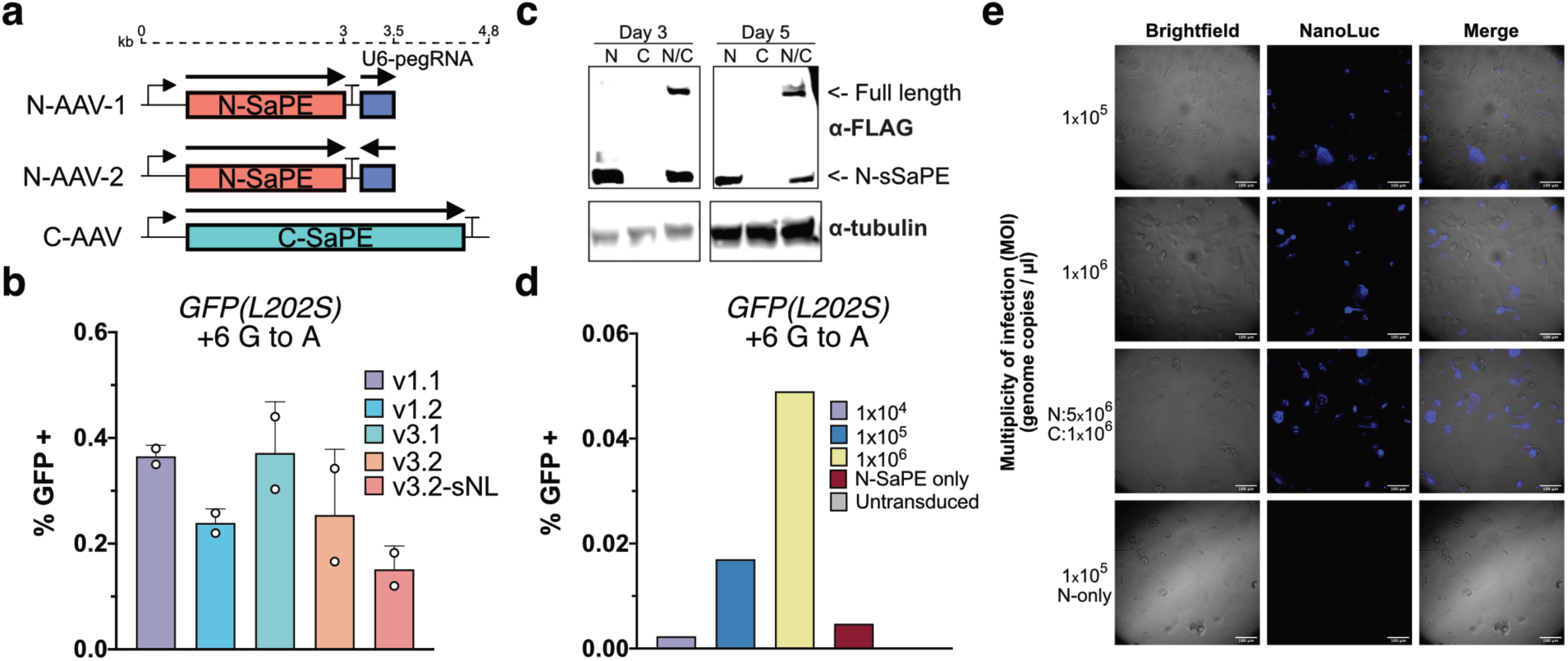
Packaging of split SaPE into AAV. (**a**) Scheme of different layouts of sSaPE in AAV genome with indicated sizes of expression cassettes. (**b**) Plasmid-based prime editing of SaPE encoded in AAV plasmids. Version 3.2-sNL corresponds to sNanoLuc replacing Npu intein. (**c**) Western blot of HEK-293T cells transduced with the indicated virus(es) for 3 or 5 days. Only the N-terminal fragment is FLAG labeled. (**d**) Co-transduction of AAVs expressing sSaPEv3 at the indicated multiplicity of infections (genome copies / cell) targeting *GFP(L202S)*. (**e**) Bioluminescence microscopy images of co-transduced AAV-sSaPEv3-sNanoLuc in U2-OS cells after 4 days.

## Discussion

We have demonstrated that an *S. aureus* prime editor presents a platform upon which in vivo prime editing can occur. While the editing rates we achieved in vitro were overall low (typically 0.5 to 1%) and not likely to be effective for many diseases, this approach offers a starting point to further refine and improve. It is important to note that comparably poor prime editing rates, albeit in plants and protoplasts, were seen in other published reports with SpPE^26,33^. Multiple engineering approaches can be undertaken to increase the efficiency of the system. To enhance activity with SaCas9, directed evolution of MMLV-RT could be employed to increase activity on pegRNA:R-loop DNA in a method akin to a recently evolved adenosine base editor^36^. A circularly permuted version of SaCas9 to facilitate the optimal RT fusion location could also be created, similar to what was also performed with base editors^37^, to increase access to the R-loop DNA. Another reason SaCas9 might have diminished prime editing rates could be attributed to a decreased residency time on the DNA target^25^. Chemically or genetically disrupting factors involved in removing Cas9 from genomic DNA has been successful at increasing other Cas9-fused effector functions. In one such study, genetic knockdown of the histone chaperone FACT increased Cas9 residence on DNA and led to improved epigenetic marking and CRISPRi^38^. It is also interesting to note that using the sSaPE-sNanoLuc platform, we see that the construct is being expressed in a majority of the cells (**Figure 3b**). A recent study examining pegRNA design principles does provide insight into optimal pegRNA design, although how well it translates to SaPE remains to be seen^39^. Further investigation, aided by the preceding suggestions, is warranted into studying the underlying reasons behind the disconnect between expression and prime editing.

A downside of SaCas9 is the more limited reach in genomic space with a longer PAM sequence of NNGRRT. This limitation is especially relevant as mutations further away from the +1 site tended to have lower editing rates. However, iterations such as SaCas9 (KKH) have lessened the PAM specificity (PAM = NNNRRT)^40^. Such codon changes could feasibly be incorporated to SaPE to expand the targeting range. Additionally, SaPE is a less ideal system for a PE3 type system, wherein a second gRNA nicks the non-editing strand to encourage the desired editing outcome, due to the stricter PAM specificity. A less stringent PAM could aid in a broader exploration of PE3 with SaPE.

Once we are able to demonstrate in vitro AAV delivery of sSaPE, the next phase will be to test in vivo. The ultimate goal is to perform delivery using targeted delivery approaches such as HUH-AAV^41^. This could have the advantage of decreasing the necessary administered dose, potentially limiting immunogenicity, and alleviating non-target cell editing concerns. In conclusion, this work provides a baseline platform for prime editor to be delivered in vivo. Through making an *S. aureus* prime editor and combining with trans splicing intein technology, we are able to package prime editor into dual AAVs for delivery. The anticipated engineering advances that will be made with prime editing components, such as increased DNA residency time or enhanced RTs, can readily be incorporated into this platform to increase editing efficiencies and move prime editing towards the clinic.

## Methods

### Nucleotide and amino acid sequences

The following tables contain sequences for pegRNAs (**Supplementary Table 1**), sequencing primers (**Supplementary Table 2**), and proteins (**Supplementary Table 3**). All oligonucleotides were synthesized by Integrated DNA Technologies (IDT).

### Cloning and DNA assembly

All prime editor expression vectors were generated using NEBuilder HiFi DNA assembly (New England Biolabs; NEB). The pegRNA expression vectors were generated as described below from Addgene plasmid #132777 (Generously provided by David Liu). AAV expression vectors were generated using NEBuilder HiFi DNA assembly combined with standard restriction digestion of the AAV vector backbone pAAV-CAG-GFP containing standard AAV2 ITRs. The *S. pyogenes* prime editor expression vector was a gift from David Liu (Addgene #132775). *S. aureus* Cas9 was amplified from a vector courtesy of Feng Zhang (Addgene #61591). Q5 site directed mutagenesis (New England Biolabs) was used to generate the Npu (C1A) mutation. Assembly reactions were transformed into competent Stellar cells (Takara Bio). Plasmid DNA was purified either as minipreps (Qiagen) or maxipreps (Thermo Fisher). DNA concentration was quantified using a Nanodrop (Thermo Fisher) and sequenced verified by Sanger sequencing (Genewiz).

### pegRNA cloning

The pegRNAs were cloned using a protocol adapted from the Liu lab^7^. These modifications were made to incorporate the *S. aureus* gRNA scaffold sequence and to add some streamlined features in regard to vector digestion and golden gate assembly cycling conditions. A detailed protocol is provided in **Supplementary Note 1**. For *S. aureus* pegRNAs, the annealing protospacer oligonucleotides were designed as such: Forward (`CACC` …[`G`] [spacer sequence]… `GTTTT`) and reverse (TACTAAAAC …[reverse complement] [`C`]…). Add a `G` prior to spacer sequence if it doesn’t begin with a `G` (and corresponding `C` on the reverse oligonucleotide). The pegRNA 3’ extension annealing oligonucleotides were designed as such: Forward (`GAGA` …[RT template + PBS]…) and reverse (`AAAA` …[reverse complement]…). The phosphorylated scaffold oligonucleotides were: Forward (/5Phos/ agtactctggaaacagaatctactaaaacaaggcaaaatgccgtgtttatctcgtcaacttgttggc) and reverse (/5Phos/ tctcgccaacaagttgacgagataaacacggcattttgccttgttttagtagattctgtttccagag). Golden gate assembly was performed with 1 µl each of 1 µM stocks of these 3 annealed oligonucleotides, 250 ng of pU6-pegRNA-RFP acceptor (Addgene #132777, courtesy of David Liu), 1 U of BsaI-HFv2 (NEB), 1 U of T4 DNA ligase (NEB), and 1x T4 DNA ligase buffer in 10 µl total volume. Reaction conditions were as follows: 10 cycles of 5 min at 37 **°**C and 10 min at 16 **°**C followed by inactivation steps of 5 min at 55 **°**C and 5 min at 85 **°**C. 1 µl of the assembly reaction was transformed into competent Stellar cells. Non-red colonies were picked for subsequent DNA isolation.

### Cell culture

HEK-293T, U2-OS, HCT116, RPE1, and HEK-293 cells were cultured in DMEM (Corning) supplemented with 10% FBS (Gibco) and 0.5% penicillin/streptomycin (Gibco). Cells were incubated at 37°C in 5% CO_2_. Bxb1-mediated recombination was used to generate the stable, single copy GFP(L202S)-2A-mKate2 HEK-293T cell line^42^.

### Plasmid transfection

Cells were seeded in a 48-well plate at 30,000 cells per well or in a 24-well plate at 60,000 cells per well. Approximately 24 hr post-seeding, cells were transfected using Lipofectamine 2000 (Invitrogen). A 1:1 molar ratio of prime editor to pegRNA vector was used (250:83 ng in 48-well plates or 500:167 ng in 24-well plates) according to the suggested manufacturer’s protocol. For split prime editor transfections, the total amount of prime editor vector was held constant. Cells were incubated for 72 hr post-transfection for all downstream analyses.

### Analysis of reversion of GFP L202S mutation

Flow cytometry analysis was carried out on a BD Fortessa X-20 instrument at the University of Minnesota Flow Cytometry Resource. Cells were prepared by first washing the cells with PBS and detaching with Accutase (Sigma). Cells were gently pelleted, washed with ice-cold PBS, and resuspended in ice-cold PBS supplemented with 5% FBS. Data from 10,000 to 100,000 cells was collected using BD FACSDiva software and compiled using FlowJo (version 10.6). Cells were initially gated based on FSC-A and SSC-A (for live cells) and then gated on FSC-W versus FSC-H (for single cells). The presence or lack of GFP expression was then evaluated (**Supplementary Note 2**).

In separate experiments, live cell imaging was carried out on an Olympus IX83 inverted microscope equipped with an Andor iXon Ultra 888 EM-CCD. Fluorescence was provided by a Sola light engine (Lumencor). For bioluminescence imaging, a Semrock light filter FF01-460/60 was used to capture NanoLuc emission. Images were processed using Fiji (version 1.51r).

### Next generation sequencing

Genomic DNA was isolated 72 hr post-transfection using the Quick-DNA Miniprep Plus kit (Zymo Research) and eluted into 25 μl 10 mM Tris-HCl, pH 8. A 150-250 bp region encompassing each targeted locus was PCR amplified from ∼40 ng genomic DNA with ends containing partial Illumina adapter sequences using CloneAmp HiFi PCR (Takara). Reaction conditions were as follows: 98 °C for 1 min, then 30 cycles of 98 °C for 10 sec, 55 °C for 10 sec, and 72 °C for 5 sec. 1 μl of each unpurified amplicon was then carried to a second PCR reaction using KAPA HiFi Library Amp (Roche Sequencing). NEBNext Multiplex Oligos (New England Biolabs) were used to add single indexes to the amplicons. Reaction conditions were as follows: 10 cycles of 98 °C for 20 sec, 61 °C for 15 sec, and 72 °C for 15 sec. 2 μl of each common amplicon were pooled and gel purified from a 1.5% agarose gel (Nucleospin clean-up, Takara Bio), eluting in 35 μl 10 mM Tris-HCl, pH 8. Common amplicon libraries were quantified with qPCR using NEBNext Library Quant kit (New England Biolabs) and pooled to equal concentrations. Sequencing was performed with an Illumina MiSeq with 2 × 150 bp paired-end reads (Genewiz). Sequencing reads were demultiplexed and analyzed using CRISPResso2 in batch mode^43^.

### GFP to BFP editing

HEK-293 cells stably expressing GFP were plated at 200,000 cells per well 24 hr prior to transfection. 250 ng of prime editor, 100 ng of pegRNA, and 50 ng of mCherry plasmid was then transfected using Lipofectamine 3000 (Invitrogen). For PE3 experiments, 100 ng of gRNA was also added. Cells were then incubated for 72 hr and analyzed using flow cytometry, gating for mCherry and BFP positive.

### PIGA editing

*PIGA*^*-*^ cell lines (HCT116 clone #2D2 or RPE1 clone #1A10) were co-transfected with 500 ng pegRNA plasmid and 1.5 µg DNA of full-length SaPE or 750 µg each of split SaPE plasmids. 1×10^6^ cells were electroporated using the Neon Transfection System (1530 V, 10 ms, 3 pulses, 10 µL tips). Transfected cells were transferred to prewarmed media in 10 cm plates and incubated for 72 hours prior to collection for downstream analysis.

### Western blot

Lysates were collected 72 hr post-transfection from HEK-293T cells in 24-well plates using RIPA buffer containing protease inhibitors. One third volume of each lysate was electrophoresed on a 4-20% SDS-PAGE gel and transferred to a nitrocellulose blot. The blot was blocked in 5% milk in TBS-T then incubated overnight at 4 °C with 1:1000 dilutions of primary antibody. Primary antibodies used were mouse anti-FLAG M2 (F1804; Sigma) or the loading control mouse anti-β-tubulin (T8328; Thermo Fisher). Blots were washed and then incubated for 1 hr with 1:10000 goat anti-mouse IgG-HRP (62-6520; Invitrogen). Blots were imaged using chemiluminescent buffer (Perkin Elmer) on an Amersham 600 UV imager (GE Healthcare).

### AAV production

All viral vectors used in this study were generated by the University of Minnesota Viral Vector and Cloning Core (Minneapolis, MN). Briefly, AAV293 cells at 60% confluence were transfected with 600 μg of DNA (viral shuttle vector encoding the payload, helper plasmid, rep/cap plasmids at 1:1:1 ratio) using polyethylenimine. 24 hr after transfection, the media was changed, and cells were checked for fluorescent protein expression (when applicable) to confirm transfection. 72 hr after transfection, cells were detached and pelleted. Viral particles were released from producer cells by repeated freeze/thaw cycles in the presence of Benzonase (100 units). Crude lysates were cleared by centrifugation and further purified using sucrose gradients. Viral particles in the supernatant were titered using qPCR with ITR-specific primers. Kanamycin-specific primers were used to confirm the absence of plasmid DNA after Benzonase treatment.

### AAV transduction

HEK-293T cells were plated 24 hr prior to transduction at 50,000 cells per well. Cells were washed with 1x DMEM (no FBS) and AAV diluted in DMEM was added gently on top. Experiments were performed at ∼1 × 10^6^ g.c.(genome copies)/cell per virus. 1 hr after virus addition, 1 ml of D10 was added on top in each well. 24 hr post-transduction, media was aspirated and 500 µl fresh D10 was added. Cells were incubated a further 48-72 hr prior to analysis.

## Data and materials availability

All materials herein are available upon reasonable request.

## Acknowledgements

We would like to acknowledge the core facilities at the University of Minnesota who assisted with this work: University Imaging Center (UIC), University Flow Resource Center (UFRC), and the Viral Vector and Cloning Core (VVCC).

## Author Contributions

E.J.A. and A.C.Z. conceived idea and designed experiments. E.J.A., A.C.Z., B.L.R., C.B.R., A.L.L, and A.T.N. carried out experiments and analyzed data. W.R.G, D.S., and E.A.H. advised and aided in assembly of the manuscript. E.J.A. authored the manuscript with input from all authors.

