## Supplementary info for "Split *Staphylococcus aureus* prime editor for AAV delivery"

**Table of contents:**


| | | |
|---|---|---|
| Supplementary Figure 1 | Conversion of GFP to BFP | 2 |
| Supplementary Figure 2 | Optimization of SaPE using GFP(L202S) reporter system | 3 |
| Supplementary Figure 3 | Prime editing in U2-OS cells | 4 |
| Supplementary Figure 4 | AAV-mediated delivery of TdTomato and SaPE | 5 |
| Supplementary Note 1 | Cloning SaPE pegRNAs using golden gate assembly | 6-7 |
| Supplementary Note 2 | Flow cytometry gating strategy and representative plots | 8 |
| Supplementary Table 1 | pegRNA sequences | 9-10 |
| Supplementary Table 2 | Primers used for genomic DNA amplification | 11 |
| Supplementary Table 3 | SaPE Protein sequences | 12-15 |

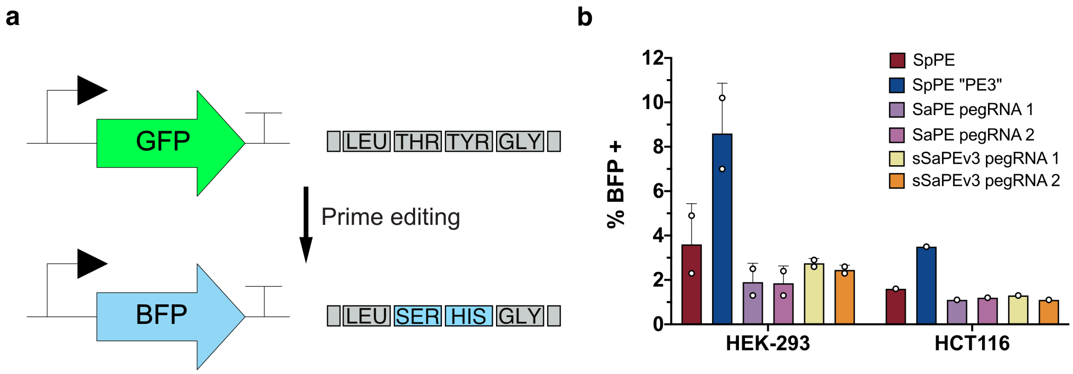


**Supplementary Figure 1 Conversion of GFP to BFP.** (**a**) Editing scheme of converting GFP to BFP through two point mutations. (**b**) Percentage of GFP converted to BFP in HEK-293 and HCT-116 cells electroporated with the indicated prime editors as analyzed by flow cytometry. Two different pegRNAs were trialed for SaPE.

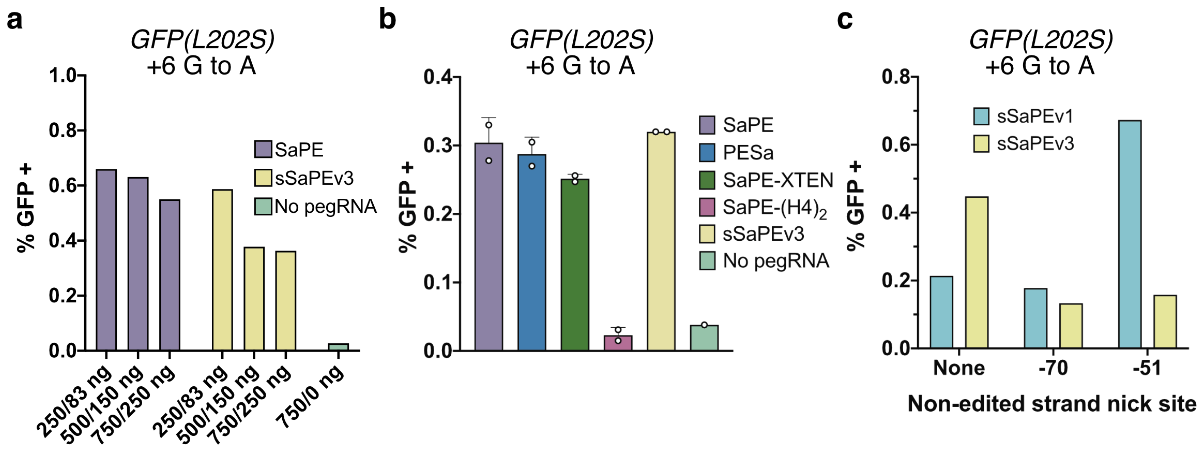


**Supplementary Figure 2 Optimization of SaPE using GFP(L202S) reporter system.** (**a**) Titration of both full-length and split SaPE. DNA concentrations on the x-axis correspond to prime editor/pegRNA amounts. (**b**) Comparison of various versions of SaPE, including altering the linker between nSaCas9 and RT. (**c**) Combining SaPE with a gRNA nicking on the non-edited strand at the indicated position to drive incorporation of the desired edit.

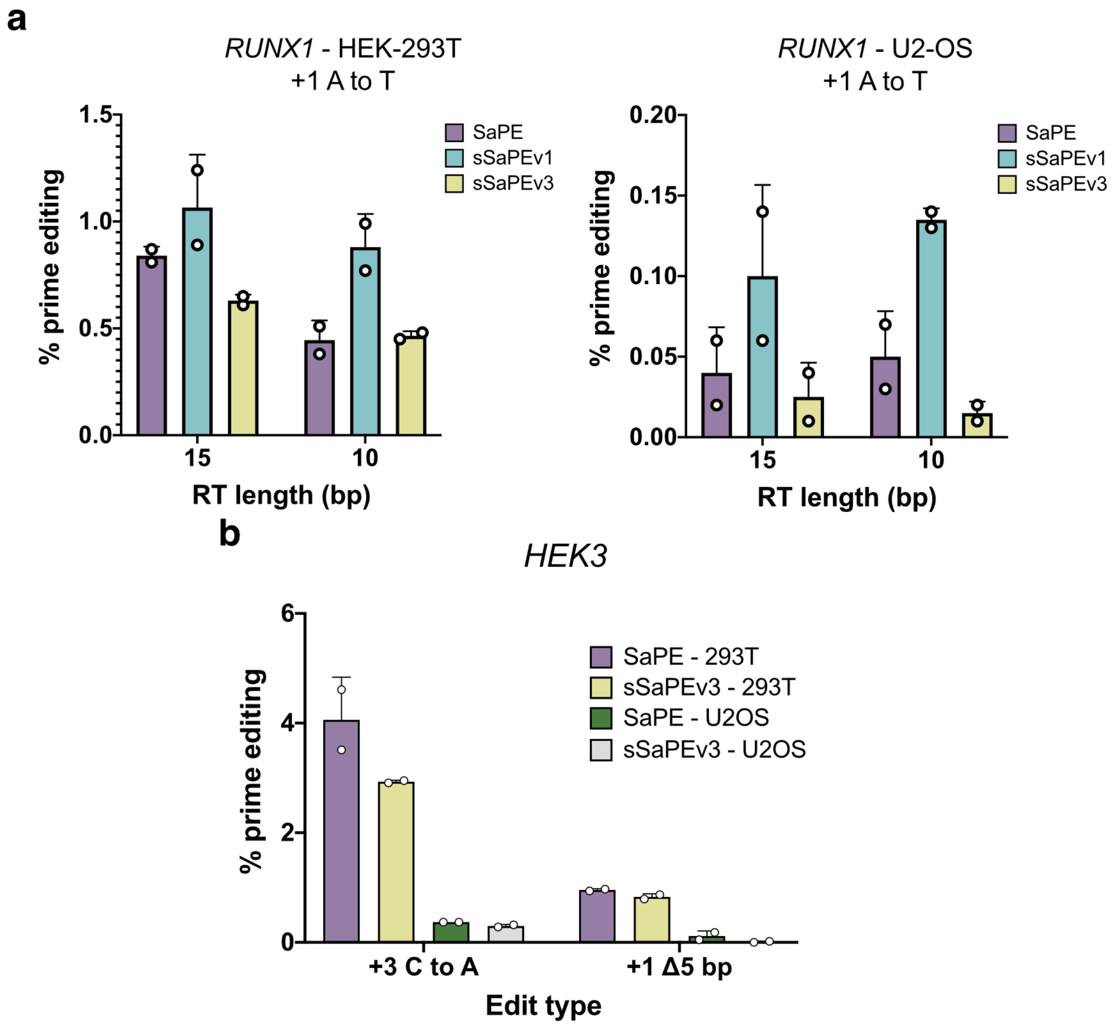


**Supplementary Figure 3 Prime editing in U2-OS cells.** (**a**) Next generation sequencing results of prime editing at the *RUNX1* locus of a +1 A to T mutation in both HEK-293T (left) and U2-OS cells (right). Different RT template lengths in the pegRNA are used while the PBS is held constant at 8 bp in length. (**b**) Prime editing comparison at the *HEK3* locus in different cell types.

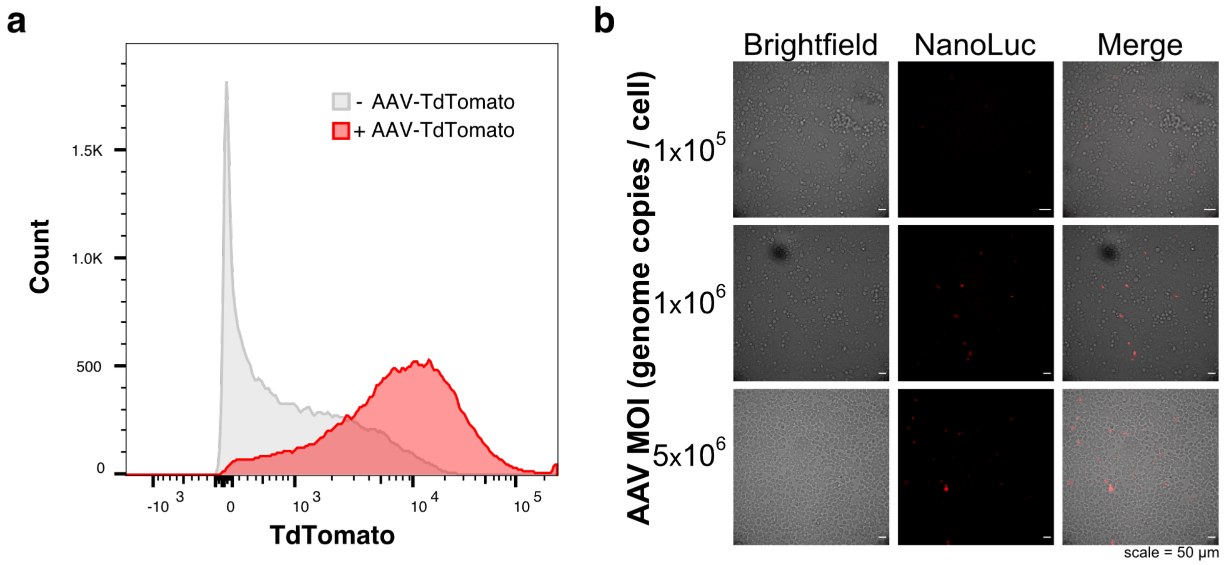


**Supplementary Figure 4 AAV-mediated delivery of TdTomato and SaPE.** (**a**) AAVs containing TdTomato as the cargo were transduced in 293T-GFP(L202S)-2A-mKate2 reporter cell line. 48 hr post-transduction, cells were analyzed using flow cytometry and compared to non-transduced cells. (**b**) Microscopic images of co-transduction of sSaPEv3-sNanoLuc encoded in AAVs in HEK-293T cells. Scale bar is 50 μm. MOI = Multiplicity of Infection.

**Supplementary note 1.** Cloning SaPE pegRNAs using golden gate assembly.

The pegRNAs were cloned using a protocol adapted from the Liu lab (Anzalone, *et al.*, 2019). These modifications were made to incorporate the *S. aureus* gRNA scaffold sequence and to add some streamlined features in regard to vector digestion and golden gate assembly cycling conditions.

**Step 0: Design of oligonucleotides**
*Protospacer oligonucleotides*
Forward: `CACC` ...[spacer sequence].... `GTTTT`
Reverse: `TACTAAAAC` …[reverse complement spacer]...
Note: Add a `G` prior to spacer sequence if it doesn't begin with a `G` (and corresponding `C` on reverse oligonucleotide).

*pegRNA 3' extension oligonucleotides*
Forward: `GAGA` …[RT template + PBS]...
Reverse: `AAAA` …[reverse complement]...

S. aureus ***phosphorylated*** *scaffold oligonucleotides*
Forward:
/5Phos/`agtactctggaaacagaatctactaaaacaaggcaaaatgccgtgtttatctcgtcaacttgttggc`

Reverse:
/5Phos/`tctcgccaacaagttgacgagataaacacggcattttgccttgttttagtagattctgtttccagag`

**Step 1: Anneal oligonucleotides**
In IDT annealing buffer (30 mM HEPES pH 7.5, 100 mM KAc), add equimolar amounts of forward and reverse oligonucleotides to 1 µM. Heat to 95 °C for 3 minutes then cool to room temperature at 0.1°C/s. Unused annealed oligonucleotide can be stored at -20°C.

**Step 2: Golden gate assembly reaction**

| | |
|---|---|
| Undigested pU6-pegRNA-RFP acceptor (Addgene #132777) | 1 µL @ 250 ng/µL |
| Annealed protospacer oligonucleotides | 1 µL @ 1 µM |
| Annealed pegRNA 3'-extension oligonucleotides | 1 µL @ 1 µM |
| Annealed **phosphorylated** sgRNA scaffold oligonucleotides | 1 µL @ 1 µM |
| BsaI-HFv2 (NEB) | 0.25 µL |
| T4 DNA ligase (NEB) | 0.50 µL |
| 10x T4 DNA ligase buffer (NEB) | 1 µL |
| $H_2O$ | 4.25 µL |
| **Total reaction volume** | **10 µL** |

Perform the following program in a thermocycler:
Cycle 10x:
5 min at 37 °C
10 min at 16 °C
Following cycles, incubate at 5 min at 55°C followed by 5 min at 85°C then hold at 12°C

**Step 3: Transformation**
Transform 1 μL of the assembly reaction into competent *E. coli* cells of your choosing. Plate on LB Agar, incubate overnight at 37 °C, and inoculate **non-red** colonies for DNA miniprep.

**Supplementary note 2.** Flow cytometry gating strategy and representative plots.

1. Gating on live cells SSC-A vs FSC-A
2. Gating on single cells (omitting aggregates) FSC-H vs FSC-W
3. Gating on eGFP negative vs positive (negative control compared to positive edit)

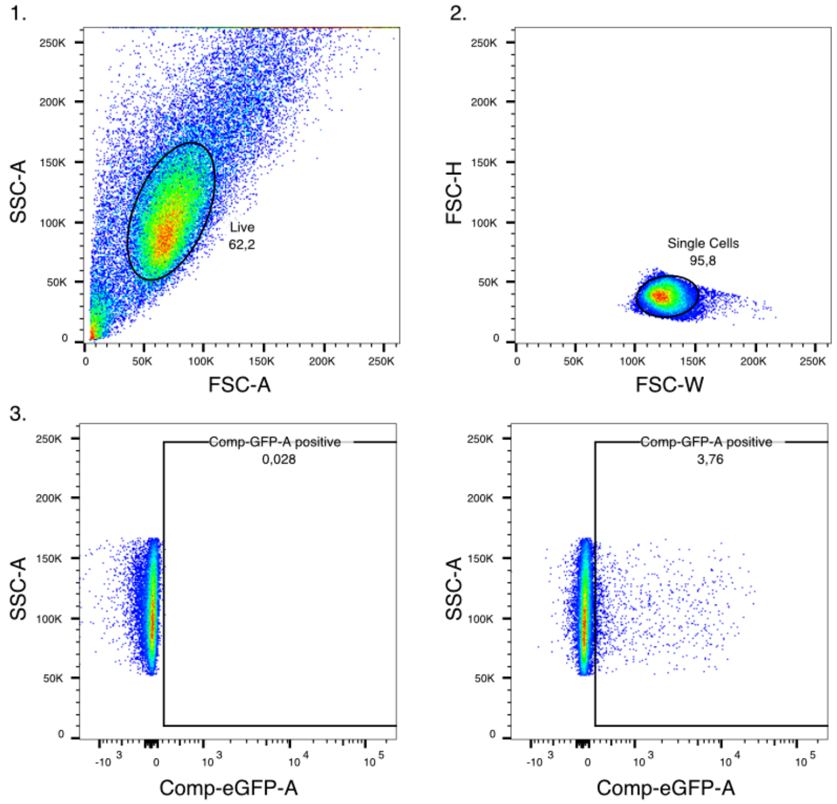

**Table 1 pegRNA sequences.**

| Target gene | Spacer sequence (5' to 3') | Modification | RT template | PBS |
|---|---|---|---|---|
| EMX1 | GGCCTCCCCAAAGCCTGGCCA | +6 G to T | ctggccactAcctgg | ccaggctttgggg |
| EMX1 | GGCCTCCCCAAAGCCTGGCCA | +6 G to T | cactAcctgg | ccaggctttg |
| EMX1 | GGCCTCCCCAAAGCCTGGCCA | +6 G to T | ctggccactAcctgg | ccaggctttg |
| EMX1 | GGCCTCCCCAAAGCCTGGCCA | +6 G to T | cactAcctgg | ccaggctt |
| L202S-GFP | TTGCTCAGGGCCGACTGGGTAC | +6 G to A | ACCACTAttTaAGtA | CCCAGTCgGCCCT |
| L202S-GFP | TTGCTCAGGGCCGACTGGGTAC | +6 G to A | ACCACTAttTaAGtA | CCCAGTCg |
| L202S-GFP | TTGCTCAGGGCCGACTGGGTAC | +6 G to A | ttTaAGtA | CCCAGTCgGCCCT |
| L202S-GFP | TTGCTCAGGGCCGACTGGGTAC | +6 G to A | ttTaAGtA | CCCAGTCg |
| RUNX1 | GTACTCACCTCTCATGAAGCACT | +6 G to C | CTTCGTACCgACAGT | GCTTCATG |
| RUNX1 | GTACTCACCTCTCATGAAGCACT | +6 G to C | TACCgACAGT | GCTTCATG |
| RUNX1 | GTACTCACCTCTCATGAAGCACT | +7 G to A | CTTCGTACtCACAGT | GCTTCATG |
| RUNX1 | GTACTCACCTCTCATGAAGCACT | +7 G to A | TACtCACAGT | GCTTCATG |
| RUNX1 | GTACTCACCTCTCATGAAGCACT | +1 A to T | CTTCGTACCCACAGa | GCTTCATG |
| RUNX1 | GTACTCACCTCTCATGAAGCACT | +1 A to T | TACCCACAGa | GCTTCATG |
| FANCF | gtagggccttcgcgcacctca | +1 A to C | gaagggattccatgC | ggtgcgcg |
| FANCF | gtagggccttcgcgcacctca | +1 A to C | gattccatgC | ggtgcgcg |
| FANCF | gtagggccttcgcgcacctca | +2 G to C | gaagggattccatCa | ggtgcgcg |
| FANCF | gtagggccttcgcgcacctca | +2 G to C | gattccatCa | ggtgcgcg |
| FANCF | gtagggccttcgcgcacctca | +3 T to G | gaagggattccaGga | ggtgcgcg |
| FANCF | gtagggccttcgcgcacctca | +3 T to G | gattccaGga | ggtgcgcg |
| GFP/BFP | GAAGCACTGCACGCCGTAGGT | +2 G to C / -1 C to T | GACCACCCTGAgCc | ACGGCGTGCAGT |
| GFP/BFP | GAAGCACTGCACGCCGTAGGT | +2 G to C / -1 C to T | ACCCTGAgCc | ACGGCGTG |
| HEK3 | tctgcttctccagccctggc | +3 C to A | ttgacccagTcc | agggctggag |
| HEK3 | tctgcttctccagccctggc | +1 5mer deletion | ggattgaccc | agggctggag |
| HEK3 | tctgcttctccagccctggc | +1 10mer deletion | cccaaggatt | agggctggag |
| HEK3 | tctgcttctccagccctggc | +1 15mer deletion | tgggccccaa | agggctggag |
| HEK3 | tctgcttctccagccctggc | +1 20mer deletion | cagtctgggc | agggctggag |
| HEK3 | tctgcttctccagccctggc | +1 25mer deletion | gtgctcagtc | agggctggag |
| FANCF | gatgttccaatcagtacgca | +6 G to C | cggcgactGtctgc | gtactgattgga |
| FANCF | gatgttccaatcagtacgca | +1 ATG_ins | cggcgactctctgcCAT | gtactgattgga |
| FANCF | gatgttccaatcagtacgca | +3 TT_ins | cggcgactctcAAtgc | gtactgattgga |
| FANCF | gatgttccaatcagtacgca | +1 tgc del | agacggcgactctc | gtactgattgga |

| | | | | |
|---|---|---|---|---|
| FANCF | gatgttccaatcagtacgca | +8 G to T | cggcgaAtctctgc | gtactgattgga |
| DNMT3B | ggtggcactgcggctggaggt | +6 G to C | ctttaaccGccacc | tccagccgcagt |
| DNMT3B | ggtggcactgcggctggaggt | +2 AA ins | ttaacccccacTTc | tccagccgcagt |
| DNMT3B | ggtggcactgcggctggaggt | +1 ggt del | tttaaccccc | tccagccgcagt |
| DNMT3B | ggtggcactgcggctggaggt | +3 $G_5$_del | ctccgctttaaacc | tccagccgcagt |

For spacer sequences that don't begin with a G nucleotide, one was added to aid in U6-driven expression of the pegRNA. Modifications are listed relative to the +1 nick site 3 nucleotides upstream from the PAM on the sense DNA strand.

**Table 2 Primers used for genomic DNA amplification.**

| Name | Sequence (5' to 3') |
|---|---|
| SpEMX1_F | ACACTCTTTCCCTACACGACGCTCTTCCGATCT CAGCTCAGCCTGAGTGTTGA |
| SpEMX1_R | GACTGGAGTTCAGACGTGTGCTCTTCCGATCT CTCGTGGGTTTGTGGTTGC |
| L202S_F | ACACTCTTTCCCTACACGACGCTCTTCCGATCT TTCAAGATCCGCCACAACAT |
| L202S_R | GACTGGAGTTCAGACGTGTGCTCTTCCGATCT GTATAGTTCATCCATGCCGAG |
| SaEMX1_F | ACACTCTTTCCCTACACGACGCTCTTCCGATCT gcaaccacaaacccacgag |
| SaEMX1_R | GACTGGAGTTCAGACGTGTGCTCTTCCGATCT agacacggagagcagctg |
| RUNX1_F | ACACTCTTTCCCTACACGACGCTCTTCCGATCT GAGGGTGCATTTTCAGGAGG |
| RUNX1_R | GACTGGAGTTCAGACGTGTGCTCTTCCGATCT CAAGCTGCCATTTCATTACAGG |
| FANCF_F | ACACTCTTTCCCTACACGACGCTCTTCCGATCT ccagagtcaaggaacacgga |
| FANCF_R | GACTGGAGTTCAGACGTGTGCTCTTCCGATCT acgtaggtagtgcttgagacc |
| DNMT3B_F | ACACTCTTTCCCTACACGACGCTCTTCCGATCT gaacccaggtagccagagac |
| DNMT3B_R | GACTGGAGTTCAGACGTGTGCTCTTCCGATCT tcctttcaacccgaacggag |
| HEK3_F | ACACTCTTTCCCTACACGACGCTCTTCCGATCT ggaaacgcccatgcaattag |
| HEK3_R | GACTGGAGTTCAGACGTGTGCTCTTCCGATCT cccagccaaacttgtcaacc |

**Table 3 SaPE Protein sequences.**

| Protein | Sequence (N-to-C) |
|---|---|
| SaPE<br>(SaCas9<br>H840A -<br>linker -<br>eMMLV-RT) | MKRTADGSEFESPKKKRKVKRNYILGLDIGITSVGYGIIDYETRDVIDAGVRLFK<br>EANVENNEGRRSKRGARRLKRRRRHRIQRVKKLLFDYNLLTDHSELSGINPYE<br>ARVKGLSQKLSEEEFSAALLHLAKRRGVHNVNEVEEDTGNELSTKEQISRNSK<br>ALEEKYVAELQLERLKKDGEVRGSINRFKTSDYVKEAKQLLKVQKAYHQLDQS<br>FIDTYIDLLETRRTYYEGPGEGSPFGWKDIKEWYEMLMGHCTYFPEELRSVKY<br>AYNADLYNALNDLNNLVITRDENEKLEYYEKFQIIENVFKQKKKPTLKQIAKEILV<br>NEEDIKGYRVTSTGKPEFTNLKVYHDIKDITARKEIIENAELLDQIAKILTIYQSSE<br>DIQEELTNLNSELTQEEIEQISNLKGYTGTHNLSLKAINLILDELWHTNDNQIAIFN<br>RLKLVPKKVDLSQQKEIPTTLVDDFILSPVVKRSFIQSIKVINAIIKKYGLPNDIIIEL<br>AREKNSKDAQKMINEMQKRNRQTNERIEEIIRTTGKENAKYLIEKIKLHDMQEG<br>KCLYSLEAIPLEDLLNNPFNYEVDHIIPRSVSFDNSFNNKVLVKQEEASKKGNRT<br>PFQYLSSSDSKISYETFKKHILNLAKGKGRISKTKKEYLLEERDINRFSVQKDFIN<br>RNLVDTRYATRGLMNLLRSYFRVNNLDVKVKSINGGFTSFLRRKWKFKKERNK<br>GYKHHAEDALIIANADFIFKEWKKLDKAKKVMENQMFEEKQAESMPEIETEQEY<br>KEIFITPHQIKHIKDFKDYKYSHRVDKKPNRELINDTLYSTRKDDKGNTLIVNNLN<br>GLYDKDNDKLKKLINKSPEKLLMYHHDPQTYQKLKLIMEQYGDEKNPLYKYYEE<br>TGNYLTKYSKKDNGPVIKKIKYYGNKLNAHLDITDDYPNSRNKVVKLSLKPYRF<br>DVYLDNGVYKFVTVKNLDVIKKENYYEVNSKCYEEAKKLKKISNQAEFIASFYNN<br>DLIKINGELYRVIGVNNDLLNRIEVNMIDITYREYLENMNDKRPPRIIKTIASKTQSI<br>KKYSTDILGNLYEVKSKKHPQIIKKGSGGSSGGSSGSETPGTSESATPESSGGS<br>SGGSSTLNIEDEYRLHETSKEPDVSLGSTWLSDFPQAWAETGGMGLAVRQAP<br>LIIPLKATSTPVSIKQYPMSQEARLGIKPHIQRLLDQGILVPCQSPWNTPLLPVKK<br>PGTNDYRPVQDLREVNKRVEDIHPTVPNPYNLLSGLPPSHQWYTVLDLKDAFF<br>CLRLHPTSQPLFAFEWRDPEMGISGQLTWTRLPQGFKNSPTLFNEALHRDLAD<br>FRIQHPDLILLQYVDDLLLAATSELDCQQGTRALLQTLGNLGYRASAKKAQICQK<br>QVKYLGYLLKEGQRWLTEARKETVMGQPTPKTPRQLREFLGKAGFCRLFIPGF<br>AEMAAPLYPLTKPGTLFNWGPDQQKAYQEIKQALLTAPALGLPDLTKPFELFVD<br>EKQGYAKGVLTQKLGPWRRPVAYLSKKLDPVAAGWPPCLRMVAAIAVLTKDA<br>GKLTMGQPLVILAPHAVEALVKQPPDRWLSNARMTHYQALLLDTDRVQFGPVV<br>ALNPATLLPLPEEGLQHNCLDILAEAHGTRPDLTDQPLPDADHTWYTDGSSLLQ<br>EGQRKAGAAVTTETEVIWAKALPAGTSAQRAELIALTQALKMAEGKKLNVYTDS<br>RYAFATAHIHGEIYRRRGWLTSEGKEIKNKDEILALLKALFLPKRLSIIHCPGHQK<br>GHSAEARGNRMADQAARKAAITETPDTSTLLIENSSPSGGSKRTADGSEFEPK<br>KKRKV* |
| FLAG-N-<br>split SaPE-<br>N-Npu<br>version 1 | MDYKDHDGDYKDHDIDYKDDDDKMAPKKKRKVGIHGVPAAKRNYILGLDIGITS<br>VGYGIIDYETRDVIDAGVRLFKEANVENNEGRRSKRGARRLKRRRRHRIQRVK<br>KLLFDYNLLTDHSELSGINPYEARVKGLSQKLSEEEFSAALLHLAKRRGVHNVN<br>EVEEDTGNELSTKEQISRNSKALEEKYVAELQLERLKKDGEVRGSINRFKTSDY<br>VKEAKQLLKVQKAYHQLDQSFIDTYIDLLETRRTYYEGPGEGSPFGWKDIKEW<br>YEMLMGHCTYFPEELRSVKYAYNADLYNALNDLNNLVITRDENEKLEYYEKFQII<br>ENVFKQKKKPTLKQIAKEILVNEEDIKGYRVTSTGKPEFTNLKVYHDIKDITARKE<br>IIENAELLDQIAKILTIYQSSEDIQEELTNLNSELTQEEIEQISNLKGYTGTHNLSLK<br>AINLILDELWHTNDNQIAIFNRLKLVPKKVDLSQQKEIPTTLVDDFILSPVVKRSFI<br>QSIKVINAIIKKYGLPNDIIIELAREKNSKDAQKMINEMQKRNRQTNERIEEIIRTTG<br>KENAKYLIEKIKLHDMQEGKCLYSLEAIPLEDLLNNPFNYEVDHIIPRSVSFDNSF |

| | |
|---|---|
| | NNKVLVKQEEASKKGNRTPFQYLSSSDSKISYETFKKHILNLAKGKGRISKTKKE<br>YLLEERDINRFSVQKDFINRNLVDTRYATRGLMNLLRSYFRVNNLDVKVKSING<br>GFTSFLRRKWKFKKERNKGYKHHAEDALIIANADFIFKEWKKLDKAKKVMENQ<br>MFEEKQAECLSYETEILTVEYGLLPIGKIVEKRIECTVYSVDNNGNIYTQPVAQW<br>HDRGEQEVFEYCLEDGSLIRATKDHKFMTVDGQMLPIDEIFERELDLMRVDNLP<br>N* |
| C-Npu-C-SaPE version 1 | MIKIATRKYLGKQNVYDIGVERDHNFALKNGFIASNSMPEIETEQEYKEIFITPHQ<br>IKHIKDFKDYKYSHRVDKKPNRELINDTLYSTRKDDKGNTLIVNNLNGLYDKDND<br>KLKKLINKSPEKLLMYHHDPQTYQKLKLIMEQYGDEKNPLYKYYEETGNYLTKY<br>SKKDNGPVIKKIKYYGNKLNAHLDITDDYPNSRNKVVKLSLKPYRFDVYLDNGV<br>YKFVTVKNLDVIKKENYYEVNSKCYEEAKKLKKISNQAEFIASFYNNDLIKINGEL<br>YRVIGVNNDLLNRIEVNMIDITYREYLENMNDKRPPRIIKTIASKTQSIKKYSTDIL<br>GNLYEVKSKKHPQIIKKGSGGSSGGSSGSETPGTSESATPESSGGSSGGSSTL<br>NIEDEYRLHETSKEPDVSLGSTWLSDFPQAWAETGGMGLAVRQAPLIIPLKATS<br>TPVSIKQYPMSQEARLGIKPHIQRLLDQGILVPCQSPWNTPLLPVKKPGTNDYR<br>PVQDLREVNKRVEDIHPTVPNPYNLLSGLPPSHQWYTVLDLKDAFFCLRLHPT<br>SQPLFAFEWRDPEMGISGQLTWTRLPQGFKNSPTLFNEALHRDLADFRIQHPD<br>LILLQYVDDLLLAATSELDCQQGTRALLQTLGNLGYRASAKKAQICQKQVKYLG<br>YLLKEGQRWLTEARKETVMGQPTPKTPRQLREFLGKAGFCRLFIPGFAEMAAP<br>LYPLTKPGTLFNWGPDQQKAYQEIKQALLTAPALGLPDLTKPFELFVDEKQGYA<br>KGVLTQKLGPWRRPVAYLSKKLDPVAAGWPPCLRMVAAIAVLTKDAGKLTMG<br>QPLVILAPHAVEALVKQPPDRWLSNARMTHYQALLLDTDRVQFGPVVALNPAT<br>LLPLPEEGLQHNCLDILAEAHGTRPDLTDQPLPDADHTWYTDGSSLLQEGQRK<br>AGAAVTTETEVIWAKALPAGTSAQRAELIALTQALKMAEGKKLNVYTDSRYAFA<br>TAHIHGEIYRRRGWLTSEGKEIKNKDEILALLKALFLPKRLSIIHCPGHQKGHSAE<br>ARGNRMADQAARKAAITETPDTSTLLIENSSPSGGSKRTADGSEFEPKKKRKV* |
| FLAG-N-split SaPE-N-Npu version 3 | MDYKDHDGDYKDHDIDYKDDDDKMAPKKKRKVGIHGVPAAKRNYILGLDIGITS<br>VGYGIIDYETRDVIDAGVRLFKEANVENNEGRRSKRGARRLKRRRRHRIQRVK<br>KLLFDYNLLTDHSELSGINPYEARVKGLSQKLSEEEFSAALLHLAKRRGVHNVN<br>EVEEDTGNELSTKEQISRNSKALEEKYVAELQLERLKKDGEVRGSINRFKTSDY<br>VKEAKQLLKVQKAYHQLDQSFIDTYIDLLETRRTYYEGPGEGSPFGWKDIKEW<br>YEMLMGHCTYFPEELRSVKYAYNADLYNALNDLNNLVITRDENEKLEYYEKFQII<br>ENVFKQKKKPTLKQIAKEILVNEEDIKGYRVTSTGKPEFTNLKVYHDIKDITARKE<br>IIENAELLDQIAKILTIYQSSEDIQEELTNLNSELTQEEIEQISNLKGYTGTHNLSLK<br>AINLILDELWHTNDNQIAIFNRLKLVPKKVDLSQQKEIPTTLVDDFILSPVVKRSFI<br>QSIKVINAIIKKYGLPNDIIIELAREKNSKDAQKMINEMQKRNRQTNERIEEIIRTTG<br>KENAKYLIEKIKLHDMQEGKCLSYETEILTVEYGLLPIGKIVEKRIECTVYSVDNN<br>GNIYTQPVAQWHDRGEQEVFEYCLEDGSLIRATKDHKFMTVDGQMLPIDEIFE<br>RELDLMRVDNLPN* |
| C-Npu-C-SaPE version 3 | MIKIATRKYLGKQNVYDIGVERDHNFALKNGFIASNCLYSLEAIPLEDLLNNPFNY<br>EVDHIIPRSVSFDNSFNNKVLVKQEEASKKGNRTPFQYLSSSDSKISYETFKKHI<br>LNLAKGKGRISKTKKEYLLEERDINRFSVQKDFINRNLVDTRYATRGLMNLLRSY<br>FRVNNLDVKVKSINGGFTSFLRRKWKFKKERNKGYKHHAEDALIIANADFIFKE<br>WKKLDKAKKVMENQMFEEKQAESMPEIETEQEYKEIFITPHQIKHIKDFKDYKY<br>SHRVDKKPNRELINDTLYSTRKDDKGNTLIVNNLNGLYDKDNDKLKKLINKSPE<br>KLLMYHHDPQTYQKLKLIMEQYGDEKNPLYKYYEETGNYLTKYSKKDNGPVIK<br>KIKYYGNKLNAHLDITDDYPNSRNKVVKLSLKPYRFDVYLDNGVYKFVTVKNLD |

| | |
|---|---|
| | VIKKENYYEVNSKCYEEAKKLKKISNQAEFIASFYNNDLIKINGELYRVIGVNNDL LNRIEVNMIDITYREYLENMNDKRPPRIIKTIASKTQSIKKYSTDILGNLYEVKSKK HPQIIKKGSGGSSGGSSGSETPGTSESATPESSGGSSGGSSTLNIEDEYRLHE TSKEPDVSLGSTWLSDFPQAWAETGGMGLAVRQAPLIIPLKATSTPVSIKQYPM SQEARLGIKPHIQRLLDQGILVPCQSPWNTPLLPVKKPGTNDYRPVQDLREVNK RVEDIHPTVPNPYNLLSGLPPSHQWYTVLDLKDAFFCLRLHPTSQPLFAFEWR DPEMGISGQLTWTRLPQGFKNSPTLFNEALHRDLADFRIQHPDLILLQYVDDLL LAATSELDCQQGTRALLQTLGNLGYRASAKKAQICQKQVKYLGYLLKEGQRWL TEARKETVMGQPTPKTPRQLREFLGKAGFCRLFIPGFAEMAAPLYPLTKPGTLF NWGPDQQKAYQEIKQALLTAPALGLPDLTKPFELFVDEKQGYAKGVLTQKLGP WRRPVAYLSKKLDPVAAGWPPCLRMVAAIAVLTKDAGKLTMGQPLVILAPHAV EALVKQPPDRWLSNARMTHYQALLLDTDRVQFGPVVALNPATLLPLPEEGLQH NCLDILAEAHGTRPDLTDQPLPDADHTWYTDGSSLLQEGQRKAGAAVTTETEV IWAKALPAGTSAQRAELIALTQALKMAEGKKLNVYTDSRYAFATAHIHGEIYRRR GWLTSEGKEIKNKDEILALLKALFLPKRLSIIHCPGHQKGHSAEARGNRMADQA ARKAAITETPDTSTLLIENSSPSGGSKRTADGSEFEPKKKRKV* |
| N-term split NanoLuc | EDFVGDWEQTAAYNLDQVLEQGGVSSLLQNLAVSVTPIQRIVRSGENALKIDIH VIIPYEGLSADQMAQIEEVFKVVYPVDDHHFKVILPYGTLVIDGVTPNMLNYFGR PYEGIAVFDGKKITVTGTLWNGNKIIDERLITPDGSMLFRVTINS |
| C-term split NanoLuc | VSGWRLFKKIS |
